# Dysregulation of neuronal activity-dependent immediate early genes in a mouse model of Angelman syndrome

**DOI:** 10.1101/2025.04.15.648865

**Authors:** Bhaskarjyoti Giri, Sagarika Das, Sudipta Jana, Nihar Ranjan Jana

**Affiliations:** Neurobiology of Disease Laboratory, Department of Bioscience and Biotechnology, Indian Institute of Technology, Kharagpur-721302, India

**Keywords:** Ube3a, Angelman syndrome, Immediate early gene, Coactivator, HDAC2

## Abstract

Angelman syndrome (AS) is a debilitating neurodevelopmental disorder triggered by impaired function of the maternal *UBE3A* gene, which codes a protein that functions as a ubiquitin ligase and transcriptional coactivator. Ube3a maternal deficient mice replicate many key behavioral deficits associated with AS; however, the underlying molecular mechanisms remain poorly understood. Using an RT^2^ Profiler PCR array that summarizes the expression of 84 genes regulating synaptic plasticity, we identified a number of dysregulated genes in the visual cortex of AS mice brains at postnatal day 25 (P25) compared to wild-type animals. In-depth analysis revealed that various immediate early genes (IEGs), like Arc, Egr1-4, and Homer, are dramatically down-regulated in the visual cortex of AS mice with regard to wild-type controls at P25. Moreover, the dark rearing of wild-type mice considerably decreased the levels of these IEGs to nearly the same level as those found in AS mice, along with the down-regulation of Ube3a. Furthermore, a significant reduction in the expression of these IEGs can also be observed in the hippocampus of AS mice. Finally, we demonstrate that the augmented activity of HDAC2 in the AS mice brain might be connected with the down-regulation of various IEGs and other synaptic plasticity-regulating genes. These findings indicate that the deregulated expression of neural activity-dependent IEGs could be linked with abnormal synaptic plasticity and associated behavioral deficits observed in AS mice.

## Introduction

Angelman syndrome (AS) is a debilitating neurodevelopmental disorder distinguished by significant intellectual disabilities, speech impairments, motor dysfunction, and recurrent seizures. Individuals with AS exhibit distinct behavioral traits, such as easily provoked laughter, flapping hand movements, hyperactivity, a fascination with water, and sleep disturbances[1–3]. The disorder arises from four different genetic anomalies, all resulting in the loss of function of the maternal *UBE3A* gene localized in the 15q11-q13 chromosomal region[4–6]. This chromosomal region is regulated by genomic imprinting, and interestingly, the paternally inherited *UBE3A* gene is explicitly silenced in neurons through the formation of a large noncoding antisense RNA transcript known as UBE3A-ATS[7,8].

The *UBE3A* gene codes a 100 kDa protein that was initially characterized as a HECT (homologous for E6-AP C-terminus) family of E3 ubiquitin ligase in the ubiquitination cycle of the ubiquitin-proteasome system (UPS) and subsequently established to work as a transcriptional coactivator of steroid hormone receptor superfamily[9–11]. Therefore, it is reasoned that altered ubiquitin ligase and coactivator functions might be linked to the pathogenesis of AS. Mounting literature now proposes that the ubiquitin ligase activity of Ube3a is vital in regulating synaptic function and plasticity[12–14]. Several substrates of Ube3a have been characterised, and many of them have been found to control synaptic function and plasticity[15–20]. These include activity-regulated cytoskeletal associated protein (Arc), Ephexin5 (a RhoA guanine nucleotide exchange factor), small conductance calcium-activated potassium channel (SK2), and calcium and voltage-dependent big potassium channel (BK) [15,18,21,20]. Abnormal accumulation of much of these substrates in the brain of Ube3a-maternal deficient mice (model mice for AS) is implicated as an underlying cause of synaptic malfunction and behavioral anomalies. AS model mice recapitulate many of the behavioral deficits observed in AS, including cognitive and motor dysfunction, audiogenic seizures, circadian clock and sleep disturbances, and anxiety-like behaviors[22–24]. These mice also demonstrate dampening of hippocampal long-term potentiation, reduced dendritic spine densities, imbalance in excitatory/inhibitory neural circuitry, and disruption of neuronal activity-dependent synaptic plasticity[25,26,19,27,28]. Many of these abnormalities could be the result of aberrant function of various substrates of Ube3a.

Although Ube3a is recognised as a transcriptional coactivator, the possible implication of this function in the pathophysiology of AS is not well understood. Ube3a is predominantly localized in the nuclear compartment, and abnormal function of nuclear Ube3a seems very crucial in the pathogenesis of AS[29,30]. This indicates that the loss of Ube3a’s coactivator function may significantly contribute to the disease’s development. Few studies have indicated that there is defective signalling of glucocorticoid hormone receptor (GR) in the AS mice brain [23]. This impairment may be associated with heightened stress responsiveness and anxiety-like behaviors observed in these mice. Additionally, Ube3a has been shown to regulate the expression of histone deacetylase 1 and 2 (HDAC1/2), and AS mice exhibit abnormal increases in the levels of both HDAC1 and HDAC2 in their brains[31,32]. Mice with overexpression of Ube3a, which serves as a typical model for autism, also display a sex-biased effect on the transcriptional deregulations of various autism-linked genes, along with on brain connectomics[33]. This suggests the importance of the coactivator role of Ube3a in regulating synaptic plasticity. In the present investigation, we explored the possible coactivator function of Ube3a in regulating neuronal activity-dependent synaptic function and plasticity using the AS mouse model. Employing an RT^2^ Profiler PCR array of 84 synaptic plasticity-regulating genes, we identified dramatically altered expression of various synaptic plasticity-regulated genes in the visual cortex of AS mice during heightened plasticity (at their age of P25). Down-regulation of various immediate early genes (IEGs) like Arc, Egr1, and Egr3 were extensively characterized in the visual cortex (under normal and dark rearing conditions) and hippocampus of AS mice brain with wild-type controls. We further demonstrate that Ube3a is a neural activity-induced gene that regulates the expression of various IEGs by modulation of HDAC2 activity.

## Materials and Methods

### Materials

Mouse monoclonal antibodies against Egr1 (sc-515830), Ube3a (sc-16689), Arc (sc-17839), and Egr3 (sc-390936) were purchased from Santa Cruz Biotechnology. The high capacity cDNA reverse transcription Kit (4368814), mouse monoclonal anti-β-actin (A5316), Trizol reagent, BCA protein estimation kit (23227), and PowerUp™ SYBR™ Green master mix (A25742) for qPCR were obtained from Sigma-Aldrich. Immobilon Western Chemiluminescent HRP substrate was purchased from Merck. Peroxidase-conjugated secondary antibodies were acquired from Vector Laboratories, whereas fluorescent-tagged secondary antibodies were sourced from Thermo Fisher Scientific. RT^2^ first strand kit (cat no 330401) and RT^2^ Profiler PCR array of mouse synaptic plasticity (330231) were purchased from Qiagen.

### Animal experimentation

The Ube3a-maternal knockout mice (AS mice) were procured from the Jackson Laboratory (strain code: 129-Ube3atm1Alb/J) and maintained in a regular IVC cage with 12 h light/12 h dark sequence and supplied appropriate quantities of pelleted diet and water. Female AS mice were crossed with wild-type males to produce offspring and afterward genotyped using genomic DNA extracted from the mouse tail as performed earlier. In most experiments, both wild-type and AS mice were sacrificed at P9 (before eye-opening) and at P25 (after eye-opening). In the dark rearing experiments, the pups were maintained in complete darkness from P0 to P25, sacrificed at P25, genotyped, and then proceeded for further analysis. Committee for the Purpose of Control and Supervision of Experiments on Animals (CPCSEA) recommendations were carefully adopted in performing all animal experimentations and permitted by the Institutional Animal Ethics Committee (IAEC) of Indian Institute of Technology, Kharagpur (protocol number: IE-4/NJ-BS/1.19).

### RT² Profiler PCR Array

Visual cortical tissues were carefully dissected out from wild-type and AS mice brains and snap-frozen in liquid nitrogen and kept at −80°C. Total RNA was isolated from the visual cortical and hippocampal tissues using the Trizol reagent following the manufacturer’s protocol, monitored by the quality assessment and determination of RNA concentrations. A total of 500 ng of RNA was then reverse-transcribed into cDNA using the RT² First Strand cDNA Synthesis Kit (Qiagen) in accordance with to the manufacturer’s instructions. The synthesized cDNA was diluted and used in the RT² Profiler PCR array for mouse synaptic plasticity in combination with the RT² SYBR green qPCR master mixture. The β-actin was used as a reference control. The cycle threshold (Ct) values acquired from the PCR experiments were exported to an Excel file, and the results were analysed in web-based GeneGlobe data analysis center (Qiagen) using the 2^-ΔΔCt method.

### Quantitative RT-PCR analysis

Total RNA was isolated from the carefully dissected visual cortical tissues and cDNA was synthesized from the total RNA with the use of high-capacity cDNA reverse transcription kit (Qiagen). Quantitative PCR (qPCR) was conducted using SYBR green universal master mixture on the Applied Biosystems QuantStudio^TM^ 6 Flex Real-Time PCR system. PCR settings used were as follows: starting denaturation at 95°C for 10 min, thereafter 40 cycles of denaturation at 95°C for 15 sec, annealing at 60°C for 1 min, and extension at 60°C for 1 min. The 18S rRNA was used as a reference control to normalize mRNA abundance. Fold changes in the transcript level were estimated using the 2^-ΔΔCt method. Primer sequences utilised to amplify transcripts were shown in Supplementary Table S1.

### Immunoblotting experiments

Mice were sacrificed through cervical dislocation, visual cortical and hippocampal tissues were cautiously dissected out, rapidly frozen into liquid N_2_, and kept at −80°C. Equal quantities of each tissue were then homogenized in RIPA lysis buffer (50 mM Tris, pH 8.0, 150 mM NaCl, 1% NP-40, 0.5% sodium deoxycholate, 0.1% SDS, and complete protease inhibitor cocktail) and were incubated on ice for 30 minutes. Homogenized samples were centrifuged at 14,000 x g for 15 minutes, supernatants were collected, and protein concentrations in each sample were measured by the BCA technique and kept at −80°C in multiple aliquots. Protein samples were mixed with SDS-PAGE sample buffer and boiled for 5 minutes and then resolved through SDS-PAGE. The resolved proteins were next transferred onto nitrocellulose membrane using semi-dry transfer apparatus. Appropriate transfer of proteins onto the nitrocellulose membrane was assessed by Ponceau staining, membrane was then washed in TBST buffer (10 mM Tris, pH 7.6, 150 mM NaCl, and 0.1% Tween-20) and blocked with 5% non-fat skimmed milk in TBST and followed by incubation with primary antibodies for overnight at 4°C. Blots were further washed and incubated with appropriate HRP-conjugated secondary antibody followed by detection with ECL reagent. The dilutions of primary antibodies used were as follows: Ube3a at 1:5000; Arc, Egr1, and Egr3 at 1:1000, and b-actin; 1: 20000.

### Immunofluorescence staining

Wild type and AS mice at their age of P25 were anesthetized by injecting ketamine (100 mg/kg body weight) and Xylazine (10 mg/kg body weight) and then perfused transcardially initially with PBS and then by 4% PFA diluted in PBS. After perfusion, brain samples were harvested, washed, and stored in 4% PFA for 24 h, followed by transfer of the brain samples into increasing concentrations of sucrose gradients (10, 20, and 30% sucrose solution) at 4°C. The brain samples were then sectioned at a thickness of 20 µm using a freezing microtome and stored in PBS having 0.02% sodium azide at 4°C. Before continuing with immunofluorescence staining method, brain sections were first incubated in antigen unmasking solution at 70°C for 40 minutes, washed several times with PBS, and exposed to endogenous peroxidases blocking solution (10% H_2_O_2_, 10% methanol in PBS) for 15 minutes. Sections were further permeabilized with 0.3% triton X-100 for 15 min, incubated with blocking solution (3% normal goat serum and 0.3% triton X-100 in PBS) for 1 hour at room temperature. After blocking, brain sections were incubated with primary antibodies against Ube3a, Arc, Egr1, and Egr3 for 12-15 h at 4°C. All primary antibodies were used at 1:50 dilutions. Sections were washed 3-4 times with PBS and further incubated with fluorescent-labelled secondary antibodies at room temperature for 1 hour. After 3-4 rounds of washing with PBS, sections on the glass slide were mounted with coverslip and imaged using a laser scanning confocal microscope (Olympus).

### Statistical analysis

Experimental data were analysed by using GraphPad Prism software (version 8). One-way analysis of variance (ANOVA) followed by Bonferroni *post-hoc* test was performed to analyse various experimental data wherever applicable. In some experiments, data were analysed by unpaired Students *t*-test. In all experiments, values were presented as mean ± SEM with *P*<0.05 was considered statistically significant.

## Results

### The visual cortex of AS mice brains exhibits an altered expression of various synaptic functions and plasticity-maintaining genes during the peak critical period

AS mice were shown to exhibit severe defects in activity-dependent synaptic plasticity in their brains[25,28]. In order to know the molecular mechanisms behind this deficit, we analysed the altered expression profile of various synaptic plasticity regulating genes in the visual cortex of AS mice brain along with wild-type controls using an RT^2^ Profiler PCR array that can detect the expression of 84 synaptic plasticity-related genes. These genes include IEGs, and other genes crucial for long-term potentiation (LTP), long-term depression (LTD), and synaptic remodelling. Visual cortex samples were acquired from wild-type, and AS mice brains during the peak critical period (at their age of P25), total RNA was isolated and processed for RT^2^ Profiler PCR array. Figure 1A shows the fold change of expression profile of all 84 genes in the AS mouse visual cortex with regard to wild-type control. Comprehensive analysis showed that the expressions of 35 genes were increased and 27 genes were decreased in the visual cortex of AS mice with respect to the control group at a cut-off value greater than 2 (Fig.1B). Among the down-regulatory genes, IEGs like Egr1-4, Arc, Homer, and JunB were showed most dramatic alterations. On the other hand, catalytic and regulatory subunits of protein phosphatases, tissue plasminogen activators, nerve growth factor receptors, and neural cell adhesion molecules exhibit considerable up-regulation. The list of all 84 genes is provided in the Supplementary Table S2. Next, we verified the expression of some of the most prominently altered genes using quantitative RT-PCR. Figures 2A and B showed the quantitative RT-PCR study data of few most down-regulated genes in the AS mice visual cortex as compared to the wild-type control group. The expression of IEGs such as Arc, Egr1-4, Homer, JunB, and Nptx2 (neuronal pentraxin 2) were significantly decreased in AS mice’s visual cortex. We have also detected significantly decreased expression of nuclear receptor 4a1 (Nr4a1, a transcription factor and master regulator of stress-induced IEGs) and kinesin family 17 (Kif17, an activity-dependent molecular motor). We also confirmed the increased expression of the catalytic subunit of protein phosphatase (PP) 1a, 2a, and 3a. All of them showed more than a 4-fold increase in their expression in the visual cortex of AS mice brain. These findings indicate that altered expression of various IEGs and protein phosphatases could be involved in disrupting activity-dependent synaptic plasticity in AS mice brains.

**Fig.1.**
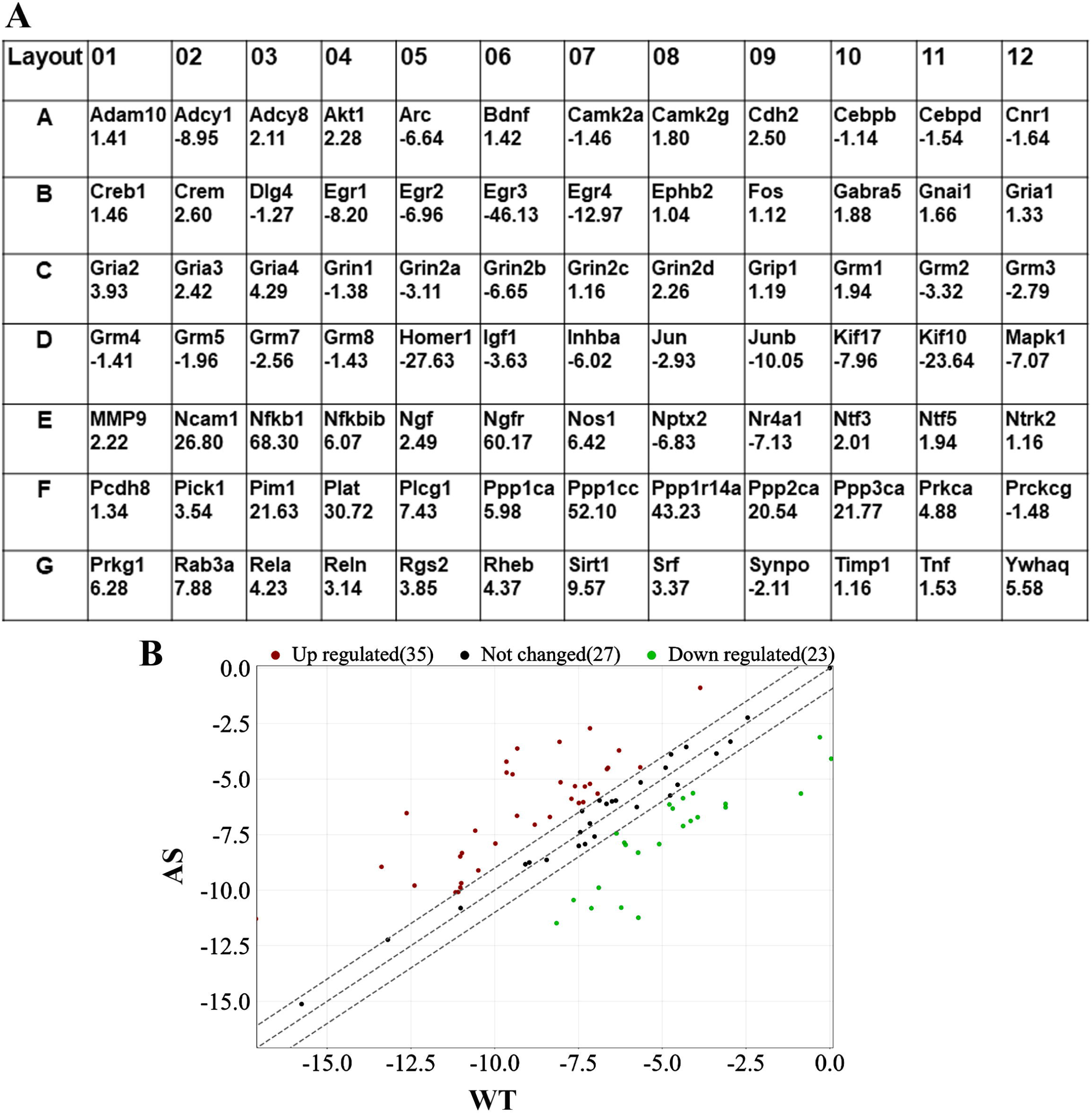
Altered expression of various synaptic plasticity regulated genes in the visual cortex of AS mice brain at P25. Visual cortical areas from wild-type (WT) and AS mice (at their age of P25) were dissected and subjected to total RNA extraction followed by RT^2^ Profiler PCR array as per the manufacturer’s protocol. Data were analysed using a web-based GeneGlobe data analysis center (Qiagen) and expressed as fold change. B) Scatter plot demonstrating genes with >2-fold difference in mRNA expression in AS mouse visual cortex at P25. In WT and AS, the log of 2^-ΔΔCt values was plotted. A detailed list (along with the full name) of the genes is provided in Supplementary Table S2.

**Fig.2.**
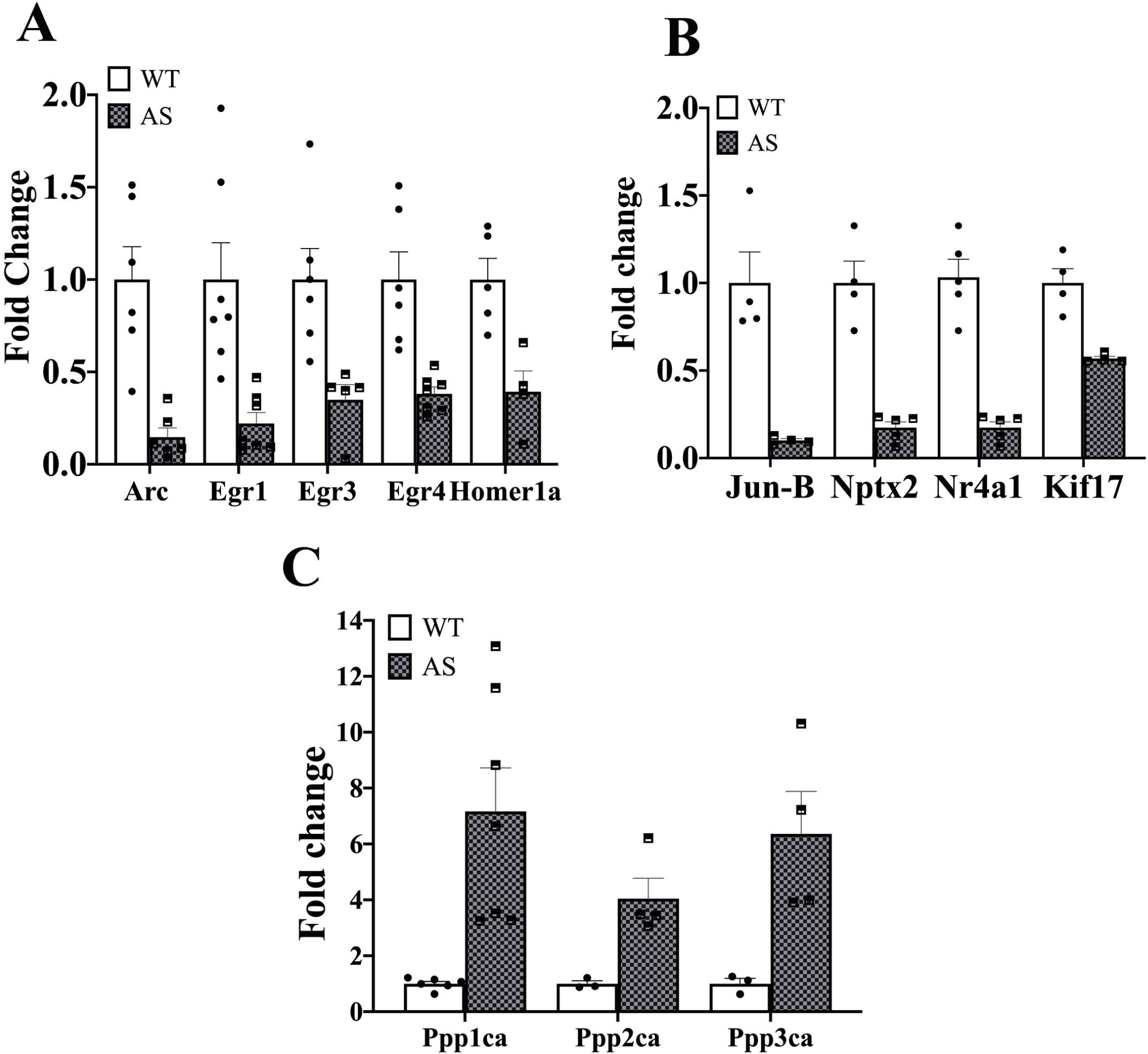
Validation of substantially altered genes in the visual cortex of AS mice through quantitative RT-PCR. Total RNA was extracted from the visual cortex of wild-type (WT) and AS mice brains as described in Figure 1, and then subjected to quantitative RT-PCR analysis of markedly up or down-regulated genes. A, B) Down-regulated IEGs and other genes. C) Up-regulated genes (catalytic subunit of protein phosphatases). Values shown are mean ± SEM with 3-6 mice samples in each experimental group. In A and B, all transcripts in AS samples are significantly down-regulated at *P*<0.001 except Kif17 (down-regulated at *P*<0.01) as compared to the WT mice group (*t*-test). In D, all the transcripts are significantly increased (*P*<0.01) when compared to the WT animal group (*t*-test). Nptx2, neuronal pentraxin 2; Nr4a1, nuclear receptor 4a1; Kif17, kinesin family 17, Ppp1ca, protein phosphatase 1 catalytic subunit α; Ppp2ca, protein phosphatase 2 catalytic subunit α; Ppp3ca, protein phosphatase 3 catalytic subunit α.

### AS mice display significant down-regulation of neural activity-dependent IEGs linked with synaptic plasticity

Next, we performed a detailed characterization of the expression and localization of various IEGs in the visual cortex of AS and wild-type mice before and after eye-opening. Visual cortex was acquired from both wild-type, and AS mice brains at the age of P9 (before eye-opening) and P25 (after eye-opening) and handled for immunoblot study of Ube3a, Arc, Egr1, and Egr3. As shown in Figure 3, Arc, Egr1, and Egr3 levels were intensely increased in wild-type mice after eye opening at P25 compared to before eye opening at P9. In AS mice, visual input also significantly increased the level of Arc, Egr1, and Egr3 at P25, but the level was considerably lower compared to wild-type controls. The level of these IEGs was decreased by nearly or more than 50% in the visual cortex of AS mice when matched to wild-type mice (Fig.3A and B). At P9, these IEGs were expressed at very low levels, and their levels were nearly similar among these two groups of animals. Remarkably, the protein and transcript level of Ube3a was dramatically up-regulated in the visual cortex of wild-type mice after eye opening at P25 compared to before eye opening at P9 (Fig.3C and D), indicating neural-activity regulated expression of Ube3a. This finding is consistent with the observations of others[15]. Next, we characterized the level and subcellular localization of Arc, Egr1, and Egr3 in the visual cortex of wild-type and AS mice brains through immunofluorescence staining. The Arc, Egr1, and Egr3 were highly expressed in the visual cortical tissues of wild-type mice brains at P25, and all of them are predominantly localized in the nuclear compartment. In the AS mice visual cortical areas, the nuclear levels of Arc, Egr1, and Egr3 were significantly decreased (Fig.4), and these results are in line with the immunoblot analysis data. We further analysed the level of Ube3a and these IEGs in the visual cortex of wild-type and AS mice in dark rearing conditions and compared them with normal rearing. Both groups of mice were raised either in normal rearing (12h light/12h dark cycle), or in dark rearing (constant darkness) conditions up to P25 and then sacrificed, and visual cortical tissues were processed for immunoblotting experiments. As shown in Figure 5, dark rearing of wild-type mice resulted in significant down-regulation of Ube3a along with Arc, Egr1, and Egr3, confirming their regulation through visual activity. In fact, the level of all three IEGs in the visual cortex of wild-type mice raised under constant darkness was comparable to that of AS mice raised under normal rearing. The dark-rearing state in AS mice showed a further decrease in the level of these IEGs when compared to normal-rearing conditions, although the decrease was not significant. From all these results, we presume that Ube3a might be directly or indirectly linked in regulating the expression of IEGs like Egr1, Egr3, and Arc.

**Fig.3.**
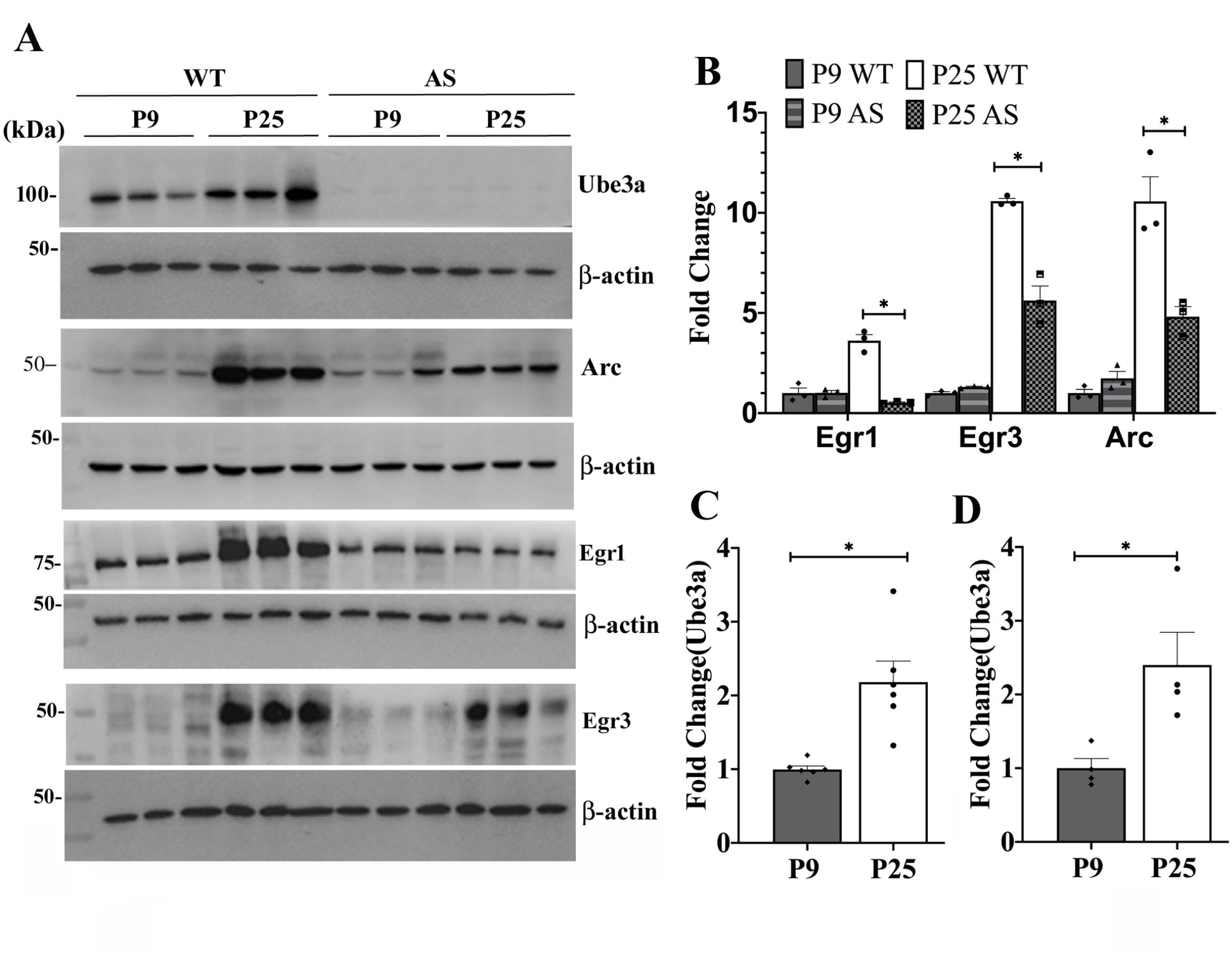
Significant down-regulation of various IEGs in the visual cortex of AS mice brain. Both WT and AS mice were sacrificed, and their age of P9 (before eye opening) and P25 (after eye-opening), visual cortex were carefully removed and processed for immunoblot analysis using antibodies against Arc, Egr1, Egr3, Ube3a, and β-actin. Band intensities of the blots were quantified, normalized against β-actin, and expressed as fold change. A) Immunoblot data. B) Decreased levels of Arc, Egr1, and Egr3 in the AS mice visual cortex at P25 compared to WT mice. Values represented are mean ± SEM with three mice samples in each group. \**P*< 0.001 compared to P25 WT group (One-way ANOVA followed by post hoc test). Each lane in the immunoblot represents a sample from different mice. C, D) Protein and mRNA levels of Ube3a were significantly increased in the visual cortex of WT mice after eye opening at P25 when compared to before eye opening at P9. C) Quantitation of immunoblot data of Ube3a shown in A. D) Quantitative RT-PCR analysis of Ube3a. In C and D, values are mean ± SEM with 4-6 animals in each group. \**P*< 0.01 compared to P9 age group (*t*-test).

**Fig.4.**
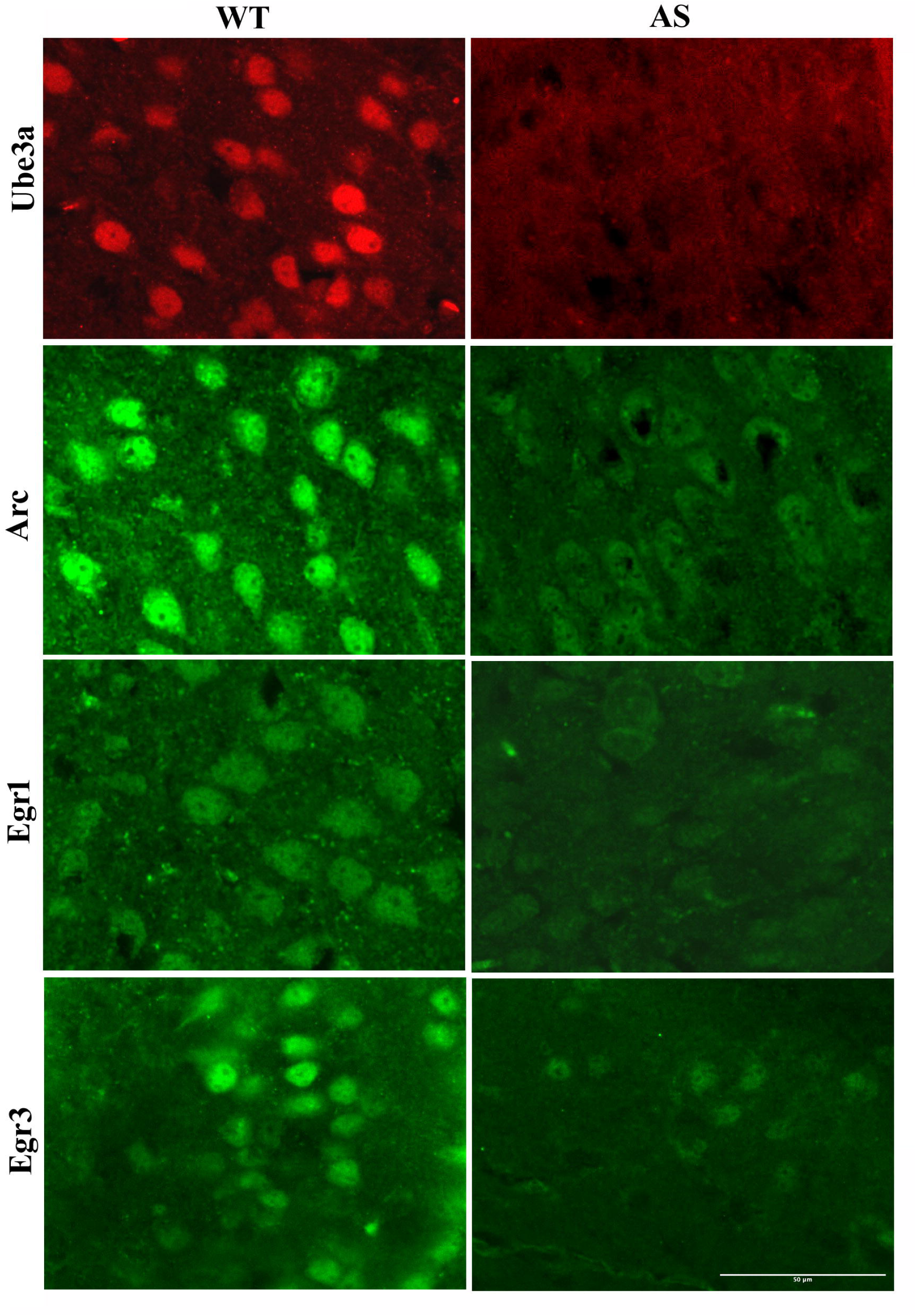
Down-regulation of Arc, Egr1, and Egr3 in the visual cortex of AS mice brain as evident from immunofluorescence staining. Brain sections with visual cortical area obtained from WT and AS mice (P25) were placed on the same slide and subjected to immunofluorescence staining with Arc, Egr1, Egr3, and Ube3a antibody followed by fluorescence-conjugated secondary antibody. Representative images of Ube3a, Arc, Egr1 and Egr3 are shown. Three mice from each group were analysed for immunofluorescence staining. Scale bar: 50 µm.

**Fig.5.**
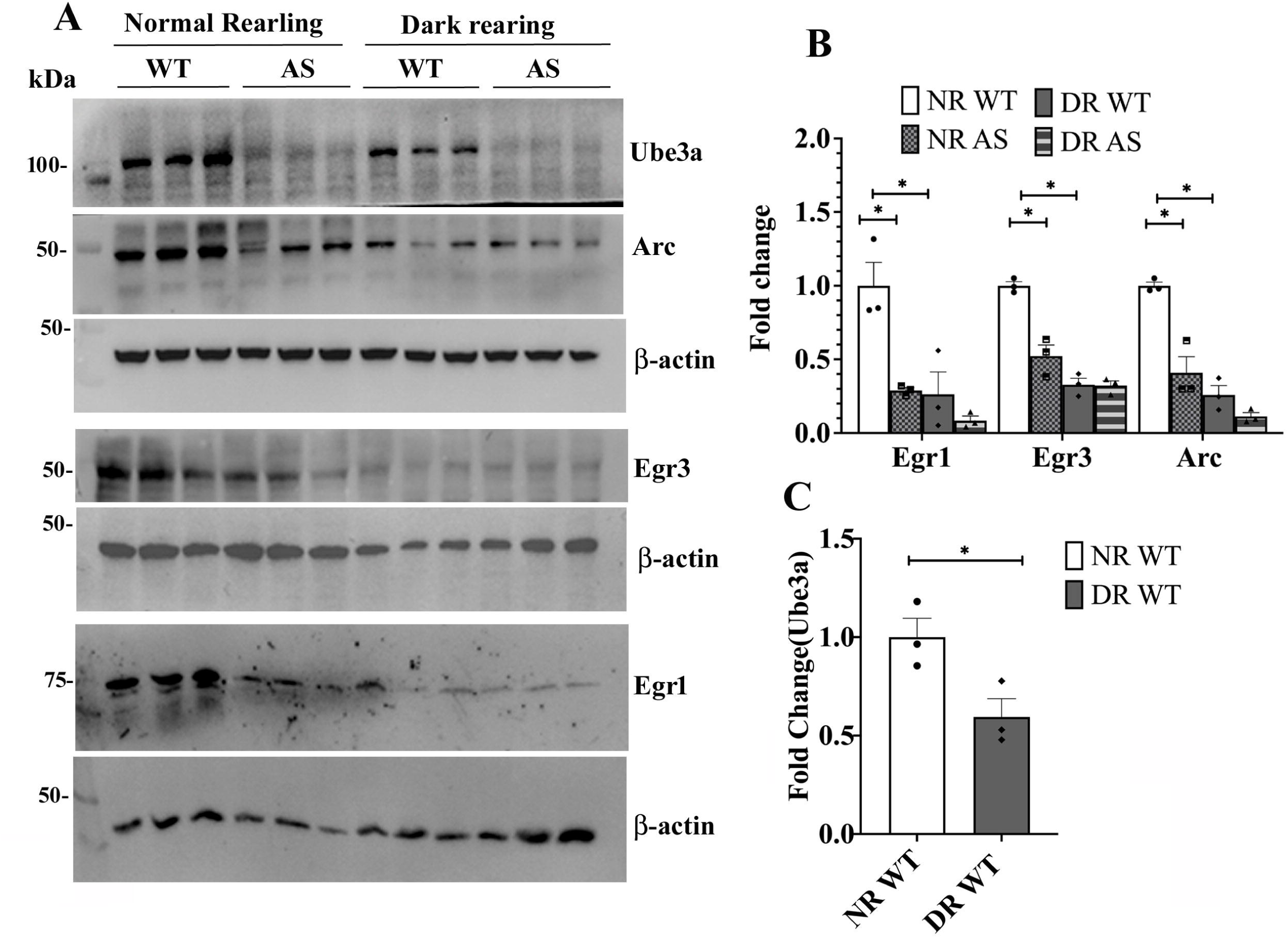
Impact of dark rearing on the expression of Ube3a and IEGs in the visual cortex of wild type and AS mice brain. Both WT and AS mice were nurtured either in the presence of normal rearing (NR, 12 h light/12 h dark cycle) or dark rearing (DR, constant darkness up to P25), sacrificed at P25, visual cortex were collected and subjected to immunoblot analysis using antibodies against Ube3a, Arc, Egr1, Egr3, and β-actin. Band intensities were quantified, normalized with β-actin, and plotted as fold change. A) Immunoblot data. B) Quantitation data of Arc, Egr1 and Egr3. C) Quantitation data of Ube3a. Values are mean ± SEM with samples from 3 animals in each group. In B, \**P*< 0.001 compared to the normal rearing NR-WT group (One-way ANOVA followed by post hoc test). Please note that the levels of Egr1, Egr3, and Arc are not significantly different between NR-AS and DR-AS groups. In C, \**P*<0.001 compared to the NR-WT group (*t*-test). Each lane in the immunoblot denotes a sample from different mice.

### Significant down-regulation of various IEGs in the hippocampus of AS mice brain

Because we observed down-regulation of various neural activity-dependent IEGs in the visual cortex of AS mice, we further checked the level of these IEGs in the hippocampus in these mice along with controls. Hippocampal tissues were obtained from the wild-type and AS mice brains at P9 and P25 and administered for immunoblot analysis with various antibodies against IEGs. Figure 6 showed that the levels of Arc and Egr3 significantly increased in the hippocampus of wild-type mice at P25 compared to P9, although the fold increase was relatively much lower than in the visual cortex. Interestingly, the Ube3a level in the hippocampus was almost similar between P9 and P25, indicating Ube3a has a different regulatory mechanism than Arc and Egr3. However, we have noticed a significantly decreased level of Arc and Egr3 in the hippocampus of AS mice at P25 brain compared to wild-type controls. This finding indicates that Ube3a regulates the expression of IEGs like Arc, Egr1, and Egr3.

**Fig.6.**
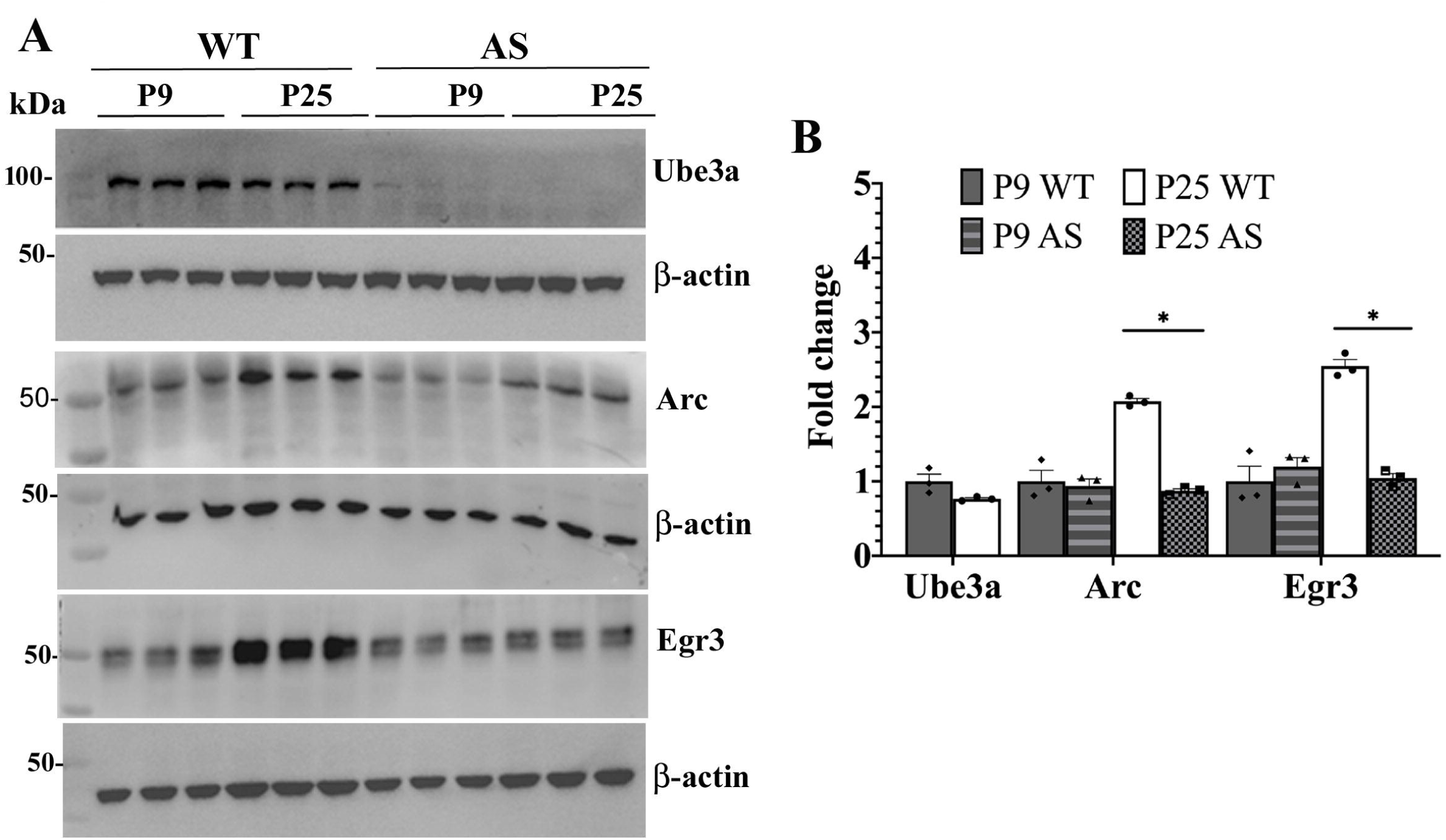
Analysis of the expression of Ube3a, Arc, and Egr3 in the hippocampus of wild-type and AS mice brains at P9 and P25. AS mice, along with WT controls, were sacrificed at their age of P9 and P25, and hippocampal tissue was collected and subjected to immunoblot analysis using antibodies against Ube3a, Arc, Egr3, and β-actin. Band intensities of each marker were quantified, normalised, and expressed as fold change. A) Immunoblot data. B) Quantitation of band intensities of Ube3a Arc and Egr3. Arc and Egr3 data were analysed by One-way of ANOVA followed by a post hoc test. Values are mean ± SEM with three different mice samples in each group. AS mice hippocampal tissues show significantly decreased levels of Arc and Egr3 (\**P*< 0.01) compared to the P25 WT group. Ube3a expression did not alter in the hippocampus of WT mice in response to eye-opening.

### Involvement of HDAC2 in Ube3a-mediated regulation of various IEGs

Since Ube3a expression is increased during neural activity and regulates the expression of various IEGs, we next explored the possible mechanism through which Ube3a could regulate the expression of these IEGs. Our earlier reports have found increased levels of HDAC1/2 in the cortical and hippocampal regions of AS mice brains as compared to wild-type controls, and HDAC2 was shown to adversely control memory formation through down-regulation of various synaptic plasticity genes, including IEGs[34,31]. Therefore, we first checked the level of HDAC2 and the acetylated histone in the visual cortex of wild-type and AS mice brains before and after eye-opening. As shown in Figure 7, the visual cortical tissues of AS mice showed a consistent increase in the level of HDAC2 together with reduced acetylated histone H3(K9) at both P9 and P25 as compared to control animals. Visual activity significantly decreased the level and activity of HDAC2 in the wild-type mice at P25. These findings show that increased activity of HDAC2 could be associated with the down-regulation of various IEGs in AS mice brains.

**Fig.7.**
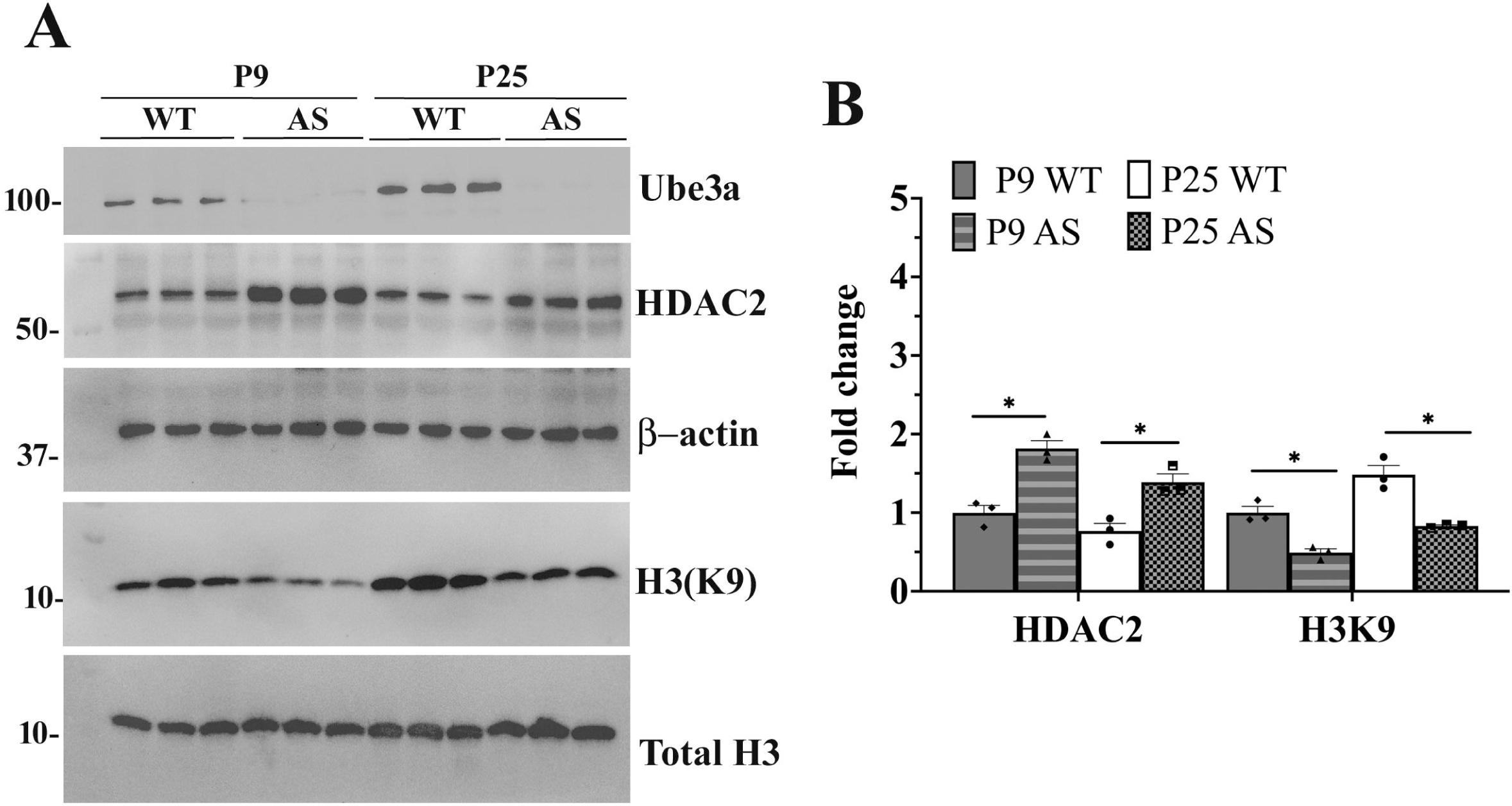
Visual cortex of AS mice brain exhibits an increased level of HDAC2 and acetylated histone H3. Visual cortical tissues from P9 and P25 animals were processed for immunoblotting experiments using various antibodies, as described in Figure 3. A) Immunoblot analysis data for Ube3a, HDAC2, total histone H3, and acetylated histone H3(K9). B) Quantitation of HDAC2 and acetylated histone H3(K9). HDAC2 data were normalised against β-actin, while acetylated histone H3(K9) was normalised against total histone H3. Values are mean ± SEM with three different samples in each group. \**P*<0.01 compared to respective age-matched wild-type control group (One-way ANOVA followed by post hoc test).

## Discussion

In the present investigation, we report altered expression of various genes linked with synapse development, transmission, and plasticity in the visual cortex of AS mice brain during peak critical period. Among the altered genes, various IEGs are found to be intensely down-regulated. Detailed characterization revealed that many IEGs like Arc, Egr1, and Egr3 are down-regulated in the visual cortex and the hippocampus of AS mice brains. Moreover, visual activity markedly regulates the Ube3a expression along with Arc, Egr1, and Egr3 in the primary visual cortical area of wild-type mice. These findings highlight that Ube3a, being a neuronal activity-regulated protein, could control the expression of various IEGs in a similar manner. We further demonstrate the possible involvement of HDAC2 in Ube3a-mediated regulation of various IEGs.

IEGs like Arc, Egr1, and Egr3 are rapidly induced in response to neuronal activity and play a crucial role in different stages of synapse development and maturation of neural circuits and their impaired function linked with many neuropsychiatric disorders[35–37]. Arc mediates endocytosis of surface AMPA receptors at the synapse and regulates long-lasting and homeostatic plasticity required for memory consolidation[38,39,37]. Arc knock-out mice display decreased numbers of mature dendritic spines in their neurons, deficiencies in experience-dependent plasticity, and deficits in learning and memory tasks[37]. Likewise, Egr1 and Egr3 deficient mice also exhibit impairment in long-term memory in various memory tests[40,36]. The EGRs are transcription factors and directly regulate the expression of Arc[41]. Therefore, our findings of down-regulation of Arc and various EGRs in the visual and hippocampal areas of AS mice brains could be connected with the impaired synaptic function and plasticity in these mice. The profound impairment of activity-dependent synaptic plasticity and maturation of the neocortex in AS mice brains could be partially due to the decreased expression of various IEGs[25,28]. Furthermore, reduced experience-mediated spine preservation shown in the visual cortex of AS mice could also be due to the down-regulation of these IEGs[27]. However, our findings do not align with other studies that showed activity-dependent Arc induction is normal in the cultured hippocampal neurons obtained from AS mice brains[42]. Although the same study reported altered homeostatic synaptic plasticity in the cultured hippocampal neuron acquired from AS mice, a feature can be seen in Arc knock-out neurons. It is also apparent that Arc is not a substrate of Ube3a[42,43]. So far, most studies have concentrated on the ubiquitin ligase activity of Ube3a in maintaining synaptic function, transmission, and plasticity and how abnormal accumulations and anomalous functions of various substrates of Ube3a ((like Ephexin5, SK2, BK) could lead to behavioral deficits in AS[18,44,17,14,20]. This study strongly indicates the involvement of the deranged coactivator role of Ube3a in aberrant synaptic function and plasticity and associated behavioral anomalies.

Another interesting result of our study is the dramatic up-regulation of catalytic subunits of PP 1a, 2a, and 3a in the visual cortex of AS mice brains. The mechanistic basis of their regulation by Ube3a and possible implications in AS pathogenesis could be interesting and require in-depth study. A recent report showed that Ube3a could regulate the activity of PP2A through the degradation of PTPA (phosphotyrosine phosphatase activator), an activator of PP2A, and increased PP2A activity can be detected in AS mice brains[19]. Pharmacological inhibition of PP2A activity in AS mice also rescued behavioral deficits[19]. However, earlier studies demonstrated increased phosphorylation of CaMKIIα at Thr286 and Thr305, along with reduced PP1/PP2A activity in the hippocampus of these mice[45]. It appears that the regulation of various protein phosphatases in AS mice brains is very complicated and requires detailed investigation.

The visual activity-dependent expression of Ube3a is not surprising as Ube3a was revealed to be regulated by neuronal activity in myocyte enhancer factor 2 (MEF2) dependent mechanisms[15]. MEF2 is a neural activity-dependent transcription factor strongly expressed in the developing brain and controls the expression of many genes (including Arc and EGRs) implicated in synaptic plasticity[46]. However, the regulation of various IEGs like Arc, Egr1-4, Homer, and Nptx2 by Ube3a is novel and seems to be mediated through the coactivator role of Ube3a. The AS mice show a consistent increase in HDAC2 activity in the visual cortical and other brain areas and HDAC inhibitors partially restore behavioral deficits in these mice. HDAC2 is known to suppress the expression of various learning and memory-related proteins, including IEGs like Arc, EGRs, and Homer and thereby, it can negatively regulate hippocampal memory formation[34]. Therefore, an aberrant increase in HDAC2 activity in the AS mice brain could be associated with the down-regulation of various IEGs. The way in which Ube3a deficiency could lead to increased HDAC2 levels in AS mice brains is not very clear. Ube3a did not interact with and control the degradation of HDAC2[32]. However, Ube3a could regulate the turnover of specific transcription factors involved in regulating HDAC2. AS mice are shown to be chronically stressed from early postnatal days, and increased corticosterone levels detected in AS mice may perhaps lead to up-regulation of HDAC2 as promoter region of HDAC2 consist GR binding site and chronic stress increased its transcription[23,47]. Moreover, steroid hormones, like estrogen and corticosterone, are also reported to induce the expression of various IEGs, including Arc, EGR1, and EGR3 (through ER/GR dependent mechanism), and Ube3a, being a coactivator of ER/GR/AR, could modulate the expression of these IEGs[48–50]. The predominant localization of Ube3a in the nucleus, abnormal function of nuclear Ube3a in AS pathogenesis, and the sex-biasing impact of Ube3a overdose on transcriptional dysregulation and autism-related behaviors strongly indicate the involvement of coactivator function in the disease pathogenesis[33,29,30].

Altogether, our findings indicate that Ube3a functions as a neural activity-dependent ubiquitin ligase that regulates the expression of various IEGs by acting as a coactivator. The dysregulation of these IEGs in the brains of AS mice may contribute, at least in part, to disruptions in synaptic function and plasticity, as well as the behavioral deficits observed in these mice.

## Acknowledgments

We would like to sincerely thank Mr. Jeet Mukhopadhyay for his technical support.

## Author contributions

All authors contributed to thestudy conception and design. Experimentations were performed by Bhaskarjyoti Giri, Sudipta Jana, and Sagarika Das; data were analysed by Bhaskarjyoti Giri and Nihar Ranjan Jana. First draft of the manuscript was written by Nihar Ranjan Jana and all authors read, commented and approved the final manuscript.

## Compliance with ethical standards

All experiments were conducted in accordance with the strict guidelines established by the Committee for the Purpose of Control and Supervision of Experiments on Animals (CPCSEA), which is part of the Ministry of Environment and Forestry, Government of India. These experiments received approval from the Institutional Animal Ethics Committee of the Indian Institute of Technology Kharagpur, under protocol number IE-4/NJ-BS/1.19.

## Data availability statement

Data will be made available by the corresponding author on request.

