## Supplementary TableS1 for "Dysregulation of neuronal activity-dependent immediate early genes in a mouse model of Angelman syndrome"

| **Gene name** | **Forward Primer** | **Reverse Primer** |
| --- | --- | --- |
| NR4a1 | 5’-CAGAGACGCGAGTGCAGC-3’ | 5’-CTCACAACATCCCCTCACCC-3’ |
| Nptx2 | 5’-AGCCAACGAGATTGTGCTGA-3’ | 5’-CCAGGTGATGCAGATGTGGT-3’ |
| Kif17 | 5’-CTATCCTCGCCCAGCAGATG-3’ | 5’-GTAGTTGGGGACAGCAGCTT-3’ |
| 18S | 5’-GAGGGAGCCTGAGAAACGG-3’ | 5’-GTCGGGAGTGGGTAATTTGC-3’ |
| JUNB | 5’-GTTTACATGGCCCCCTTCCA-3’ | 5’-GGTCCCTCCCTAGTATCCCC-3’ |
| Arc | 5’-ACCTGACATCCTGGCACCTC-3’ | 5’-GGTCCCTCCCTAGTATCCCC-3’ |
| Egr3 | 5’-GATCGGGAAGGCTTGGTTGG-3’ | 5’-TCCCAAGTAGGTCACGGTCT-3’ |
| Egr4 | 5’-CTCCACCTGAGCGACTTCTC-3’ | 5’-TCCAGGAAGCAGGAGTCTGT-3’ |
| Ppp1r14a | 5’-AGGACTTCGTCCAGGAGCTA-3’ | 5’-AACAATCTGGGCTTTGGGGG-3’ |
| Ppp1cc | 5’-GAAAAAGAAGCCCAACGCCA-3’ | 5’-GTATAAACCGGTGGACGGCA-3’ |
| Ppp2ca | 5’-AAGGTTCGTTACCGAGAGCG-3’ | 5’-GGTGACAGACCACCGTGTAG-3’ |
| Ppp3ca | 5’-GGTGGTGAAAGCCGTTCCAT-3’ | 5’-CACAAACTGTGACTGGTGCG-3’ |
| EGR1 | 5’-AGCGAACAACCCTATGAGCA-3’ | 5’-TCGTTTGGCTGGGATAACTC-3’ |
| Homer1a | 5’-CCAGAAAGTATCAATGGGACAGATG-3’ | 5’-TGCTGAATTGAATGTGTACCTATGTG-3’ |
| Ube3a | 5’-GATGAAAAGTCGTAAGGGTCAG-3’ | 5’-GTGTTTTTAGCCTCTGCTC-3’ |

**Supplementary Table S1: List of primer sets used in the study**
